# Expression screen of TNFR1^R347A^, MyD88, IRAK4 death domains in E. coli followed by purification and biophysical characterization of TNFR1^R347A^ death domain

**DOI:** 10.1101/2024.12.13.628329

**Authors:** Kamil Przytulski, Aleksandra Podkówka, Tomasz Tomczyk, Daria Gajewska, Magdalena Sypień, Agnieszka Jeleń, Priyanka Dahate, Anna Szlachcic, Michał Biśta, Michał J. Walczak

## Abstract

Death domains play a crucial role in signaling pathways related to inflammation and programmed cell death, rendering them promising targets for therapeutic interventions. However, their expression as recombinant proteins often pose challenges. Here, we present expression screening of TNFR1, IRAK4, and MyD88 death domains in *E. coli*, followed by the biophysical characterization of TNFR1 death domain after subsequent construct optimization. The study also discusses the influence of pH and ionic strength on TNFR1^R347A^ stability, providing statistical models to predict optimal conditions of the buffer to achieve the highest protein stability.

**Highlights:**

- Optimization of expression conditions for TNFR1^R347A^, MyD88, IRAK4 death domains in *E. coli* BL21(DE3) cells.
- High-yield production of soluble monomeric TNFR1^R347A^ death domain.

## Introduction

Tumor necrosis factor receptor-1 (TNFR-1) plays a crucial role in cellular signaling. It is activated by the interaction with its ligand, Tumor Necrosis Factor (TNF), initiating intricate intracellular signaling cascades. These pathways orchestrate the activation of the transcription factor NF-κB, pivotal in regulating gene expression, and initiating mechanisms leading to programmed cell death. TNF not only induces apoptosis in specific tumor cells but also serves as a key mediator in the inflammatory response. However, dysregulated TNF production or prolonged activation of its signaling pathways has been linked to the development and progression of various human diseases, including rheumatoid arthritis, inflammatory bowel disease, sepsis, cerebral malaria, diabetes, and autoimmune disorders. Therefore, blocking the TNF pathway has been established as a therapeutic strategy validated by numerous biological drugs on the market [1]. Interleukin-1 Receptor-Associated Kinase 4 (IRAK4), a member of the kinase family (including IRAK1, IRAK2, and IRAK-M), is a serine/threonine kinase integral to transduction pathways activated by Toll-like receptors (TLRs) and Interleukin-1 (IL-1) family receptors [2]. Activation of those receptors by the foreign pathogens identification or inflammatory signals detection, prompts the binding of IRAK4 to the adapter protein MyD88. This interaction triggers IRAK4’s activation, subsequently initiating the NF-κB pathway, which culminates in the production of pro-inflammatory cytokines [3]. Numerous IRAK4 inhibitors have been designed to impede the protein’s kinase activity, with the most progressed ones advancing to Phase II clinical trials. However, some studies suggest that in specific cell types, the kinase activity might not be crucial for disease relevance. Recently, a new modality has emerged that does not require a tractable catalytic site. Targeted Protein Degradation (TPD) has the potential to broaden the scope of therapeutic targets [4]. Consequently, not only blocking the IRAK4 kinase activity, but degradation of IRAK4 utilizing a PROTAC molecule [5] resulted in disrupted signaling pathways, leading to improved therapeutic outcomes [6].

Due to inherent tendency of death domains for oligomerization [7], [8], they are prone to aggregate during recombinant protein production. Our research focused on developing the general strategy and identifying the optimal conditions for producing soluble death domains. In the present work attempts have been made to optimize conditions to produce different soluble death domains derived from TNFR1, IRAK4 and MyD88 in *Escherichia coli* BL21(DE3) host cells. Various host cells, including bacteria, yeast, insects, plants, and animals serve as platforms for recombinant protein production. Since early 1980s, *Escherichia coli* has been the preferred method for expression of recombinant proteins from both eucaryotic and prokaryotic species. This preference persists due to its ability to thrive in low-cost culture media under precisely controlled laboratory conditions. Its rapid growth, with doubling time within 20- 30 minutes and ease of incorporate foreign DNA, makes the entire process of clone selection robust and convenient. For optimization we focused on key parameters affecting production of soluble protein in *E. coli* [9]. Highly potent promoters (e.g. T7) and elevated inducer concentrations often lead to substantial protein production, overwhelming the folding machinery. This results in an imbalance between the rates of protein translation and proper folding, resulting in aggregation. Properly controlling protein expression levels is crucial to mitigate this issue and ensure accurate protein folding processes and attenuate protein aggregation [10]. Elevated temperatures tend to favor protein aggregation due to the increased strength of hydrophobic interactions, which play an important role in driving the aggregation process. Conversely, lowering the cultivation temperature is a widely acknowledged strategy to mitigate the occurrence of protein aggregation within living cells. This approach minimizes the impact of hydrophobic interactions, thereby reducing the likelihood of proteins clumping together and forming aggregates [11]. Fusion proteins not only aid in the proper folding of proteins by facilitating the acquisition of their correct structure but also serve to safeguard the recombinant protein from potential degradation by intracellular proteases [12]. Here, we used the small ubiquitin-like modifier (SUMO), an 11 kDa protein, present in both yeast and vertebrates. SUMO has shown its ability to improve protein stability and solubility as an N-terminal fusion. While SUMO itself does not participate in purification, incorporating a hexahistidine tag facilitates the purification process, and afterwards SUMO domain can be specifically removed using Ulp1 protease [13], [14].

Ultimately, to confirm the feasibility of large-scale protein purification, herewith we provide a robust protocol for the production of monomeric TNFR1^R347A^ mutant. Death domains are prone to form aggregates under physiological pH conditions, which could be challenging especially in structural studies. To address this issue, relatively low (<4) or high (>8) pH are employed to mitigate the risk of self-association. The wild-type TNFR-1 DD demonstrated high insolubility across a broad pH range (pH 4-10), requiring the generation of mutant proteins to enhance solubility and stability. The mutation site R347 was strategically chosen due to its pivotal role in mediating both cytotoxicity and binding ability of TNFR-1 DD, as supported by prior investigations. Notably, the R347A mutant displayed enhanced solubility and stability under elevated pH conditions (pH 8.8), facilitating the determination of secondary structure and offering valuable insights into the protein’s topological fold [7], [15]. We characterized TNFR1 death domain mutant R347A, by investigating the impact of pH and ionic strength on protein stability and mathematically described their influence.

## Results and discussion

### Various death domains are expressed as soluble proteins in E. coli

To determine the optimal conditions to produce soluble death domains of TNFR1, IRAK4, and MyD88 proteins, we focused on several crucial factors influencing protein production. All proteins were produced in fusion with a C-terminal hexahistidine tag, allowing purification using metal-chelating chromatography matrices with capacity to capture free electron pairs [16], [17]. The study investigated three distinct culture conditions, varying in temperature and incubation time after chemical induction of the gene encoding the target protein of interest. To trigger gene expression, Isopropyl β-D-1- thiogalactopyranoside (IPTG), a non-metabolizable lactose derivative, was introduced into the medium across different concentrations.

We chose to utilize prior literature findings and explore the production of death domains in fusion with a C-terminal hexahistidine tag [7], [15]. Employing the SDS-PAGE methodology [18], we conducted a comprehensive examination of the protein composition within distinct cellular fractions (Figure 1). Specifically, our attention was directed towards both whole cell lysates (WCL) and the soluble proteins fraction. Soluble proteins were subjected to incubation with NiNTA resin to ascertain there is no potential formation of soluble aggregates, which could impair the following high-scale purification. Cellular lysates and soluble fractions were normalized based on optical density (OD). On the contrary, protein fractions bound to the chromatographic resin were normalized with respect to the culture volume. Such an approach facilitated a more accurate determination of the actual protein quantity recoverable from the particular culture volume and allowed for estimation of host cell proteins derived contamination. Cells were subjected to lysis in the presence of lysozyme (appx. 14 kDa), enhancing sample consistency. The highest OD values were observed under conditions in which cells were cultured at 25°C overnight. For the majority of samples, the addition of IPTG to the culture medium resulted in a decrease in OD of cultures after the incubation (*Supplementary*: Figure S1). In case of TNFR1 and IRAK4, the production level of soluble protein within a given amount of cells was similar across all examined conditions. MyD88 death domain deviated and exhibited higher insolubility compared to TNFR1 and IRAK4. Since the OD of cultures conducted overnight at 25 °C was the highest, and considering the unified protein production within the cells, the highest production efficiency per culture volume would be anticipated for cultures conducted overnight at 25 °C. This observation was also confirmed due to analysis of eluate fractions. In summary, the optimal condition to produce death domains derived from TNFR1, IRAK4, and MyD88 encompass supplementing Luria-Bertani broth (LB) with 100 μM IPTG, followed by incubation of bacterial cultures overnight at 25 °C.

**Figure 1.**
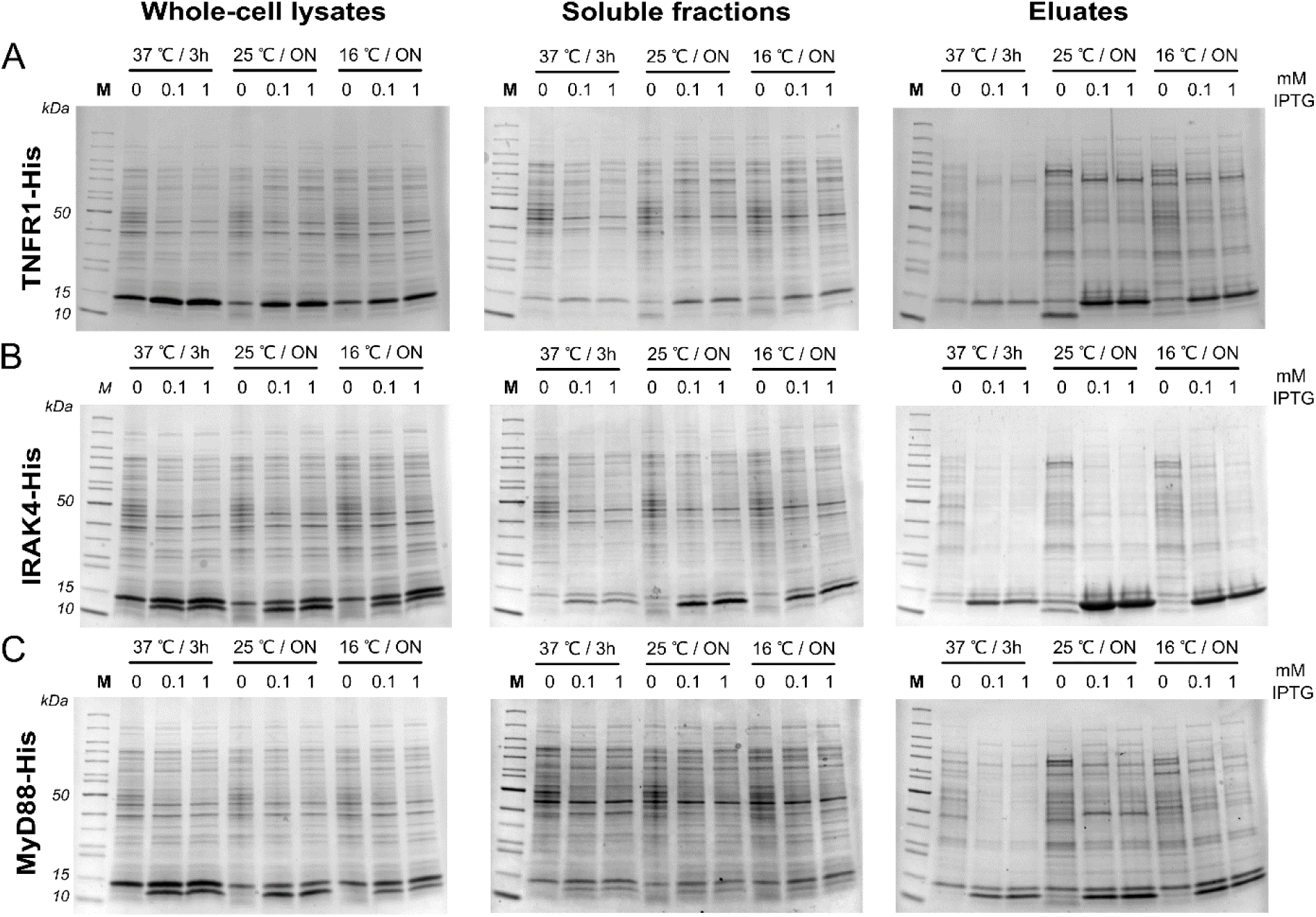
Analysis of TNFR1 **(A)**, IRAK4 **(B)** and MyD88 **(C)** expression in E. coli BL21(DE3) strain. Death domains were expressed in fusion with C-terminal hexahistidine tag. Protein production was induced either with 0.1 or 1 mM Isopropyl β-D-1-thiogalactopyranoside (IPTG). Following induction, bacterial cultures were subjected to incubation under distinct conditions: (I) 37 ℃ / 3 hours, (II) 25 ℃ / overnight (ON), and (III) 16 ℃ / ON. Normalization of whole cell lysates, and soluble fractions were performed based on OD_600_, with a loading quantity of 20/OD. After incubation of soluble fraction with Ni-agarose beads, bound proteins (eluate fractions) were normalized, loading cells derived from 0.2 mL of culture. MW: TNFR1-His – 13.8 kDa; IRAK4-His – 12.79 kDa, MyD88-His – 12.62 kDa; Lysozyme – appx. 14 kDa

### TNFR1^R347A^ construct optimization

The production yield of soluble TNFR1 death domain was relatively low, prompting us to undertake further optimization. For this purpose, we employed a similar scheme of conditions as described in the previous paragraph. However, this time we focused on testing various TNFR1 constructs (Figure 2D) that would enable enzymatic cleavage of the hexahistidine tag. Positioning of fusion partners at the N-terminus or C-terminus of the passenger protein can induce distinct effects. N-terminal tags exhibit superiority over their C-terminal counterparts due to the following reasons: (I) they establish a reliable context for efficient translation initiation by capitalizing on translation initiation sites on the tag; (II) upon removal, they yield minimal or negligible additional residues at the native N-terminal sequence of the target protein, as the majority of proteases cleave at or proximal to the C-terminus of their recognition sites [19].

**Figure 2.**
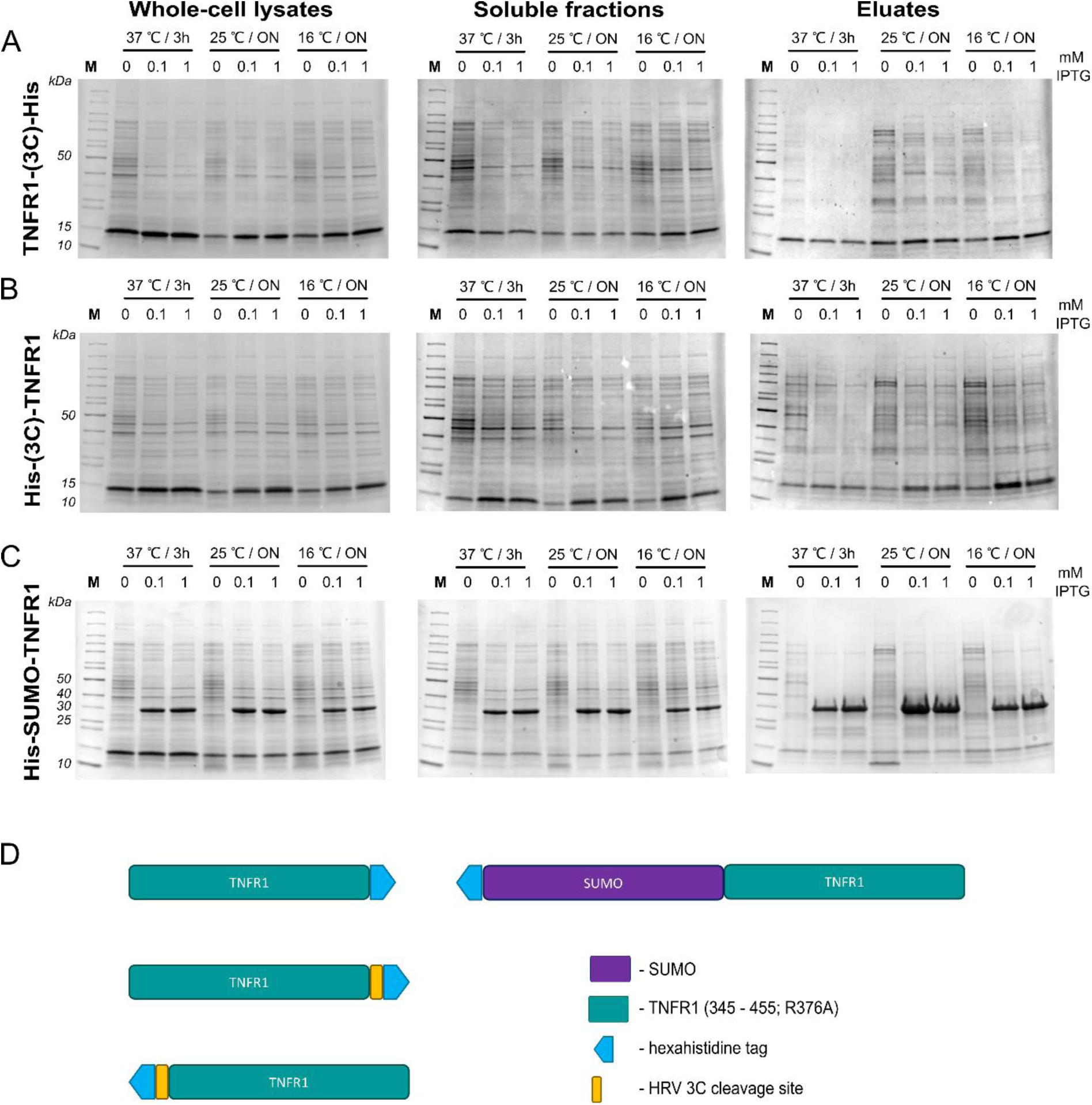
Analysis of TNFR1-His **(A)**, His-TNFR1 **(B)** and His-SUMO-TNFR1 **(C)** expression in E. coli BL21(DE3) strain. Death domains were expressed in fusion with C-terminal or N-terminal hexahistidine tag or as a His-SUMO fusion. Protein production was induced either with 0.1 or 1 mM Isopropyl β-D-1-thiogalactopyranoside (IPTG). Following induction, bacterial cultures were subjected to incubation under distinct conditions: (I) 37 ℃ / 3 hours, (II) 25 ℃ / ON, and (III) 16 ℃ / ON. Normalization of whole cell lysates, and soluble fractions were performed based on OD_600_, with a loading quantity of 20/OD. After incubation of soluble fraction with Ni-agarose beads, bound proteins (eluate fractions) were normalized, loading cells derived from 0.2 mL of culture. MW: TNFR1-(3C)- His – 14.68 kDa; His-(3C)-TNFR1 – 15.29 kDa, His-SUMO-TNFR1 – 26.14 kDa; Lysozyme – appx. 14 kDa **(D)** Schematic representation of tested TNFR1 death domain constructs.

Due to the presence of hydrophobic patches on the surface of death domains, we chose to cleave the protein at a low temperature to reduce the hydrophobic effect. Cleavage of the fusion protein was accomplished with the Human Rhinovirus (HRV) 3C protease, known for its activity at low temperatures unlike the widely used tobacco etch virus (TEV) protease [20]. To suffice high solubility, we used SUMO fusion construct. An additional benefit of SUMO fusion protein is that it can be processed by a specific protease, Ulp1, which leaves no additional amino acids on the target protein, allowing to obtain the endogenous protein sequence [13]. Based on our results, it can be observed that the production of TNFR1 in fusion with a C-terminal hexahistidine tag, with preceding HRV 3C recognition site, reduced the yield of the obtained protein, when compared to the protein produced in fusion with a non-cleavable his-tag (Figure 1A). Furthermore, protein production with cleavable His-tag performed better in the case of the construct in which tags were attached to the N-terminus, and protein production was conducted overnight at 16 °C. Ultimately, the most efficient protein production was achieved with the TNFR1- SUMO construct.

For high-scale production, we chose to induce gene expression using 0.1 mM IPTG followed by incubation of bacterial host cells overnight at 25 °C. While conditions can still be further optimized, employing DoE approach to refine values of three variables within different ranges: (I) OD: 0.4 – 0.8; (II) IPTG: 0.05 – 1 mM; (III) Temperature: 16 – 33 °C, we decided to initiate purification at this stage, to assess protein production yield.

### TNFR1^R347A^ in fusion with SUMO results in high yield of protein production

Bacterial culture described in the previous paragraph was scaled up to 2 L. Before the purification, the impact of polyethyleneimine (PEI) on protein solubility was assessed. PEI is a polycationic reagent commonly used for precipitating nucleic acids from samples [21]. If a protein interacts with nucleic acids or carries a high negative charge in low ionic strength buffers, it may lead to protein precipitation. From our experience, 1 M NaCl prevents the precipitation even of strongly negatively charged proteins (e.g. MsyB [22]), and selectively precipitates nucleic acids (data not shown). In the case of TNFR1 protein, we did not observe its precipitation in the presence of 0.1% PEI and 0.3 M NaCl (Figure 3A). For protein purification, we utilized a nickel column and did not detect significant amounts of protein in the unbound flow-through fraction. The enzymatic cleavage of the fusion protein by the Ulp1 protease was highly efficient. Following cleavage and reloading onto the nickel column, we did not observe any re-binding of TNFR1 to the chromatography matrix, indicating that endogenous histidine residues did not enable reassociation with the beads.

**Figure 3.**
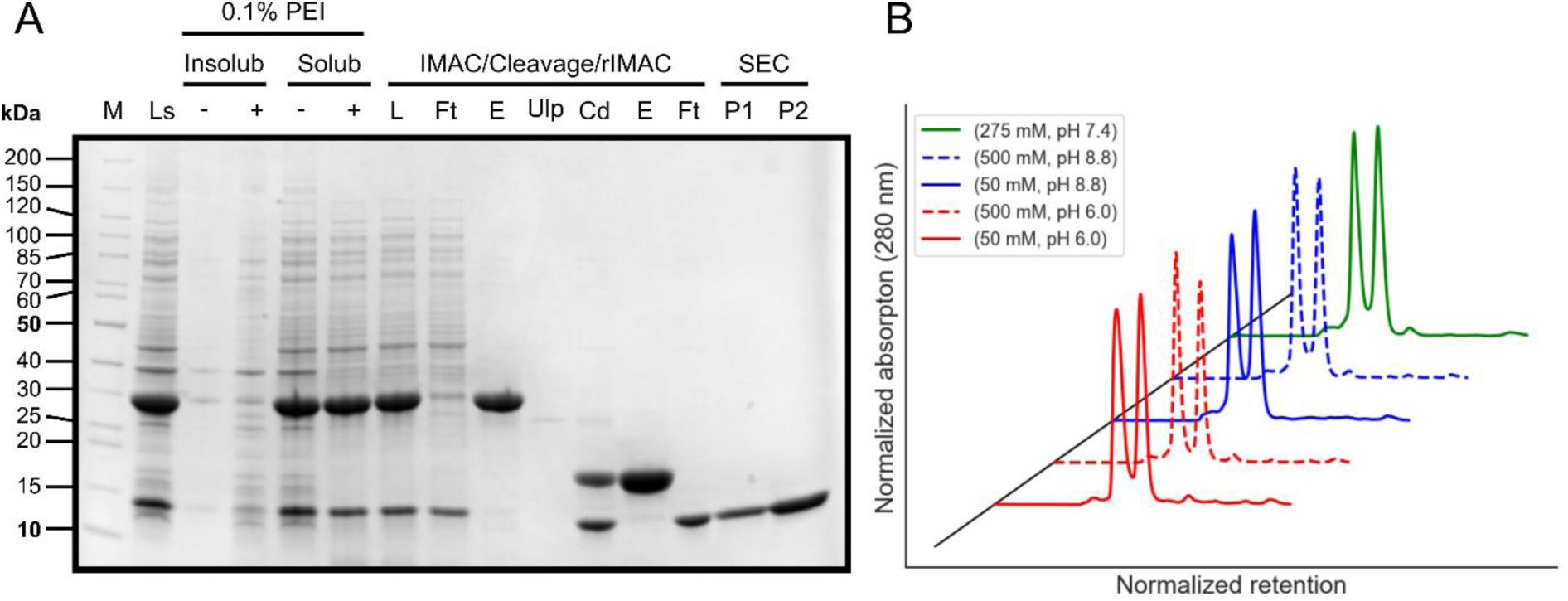
Image of the samples collected during a high-scale purification, analyzed with SDS-PAGE **(A)**. Cells lysate prior to clarification by centrifugation [**Ls**]. The effect of 0.1% polyethyleneimine [**PEI**] on protein solubility was tested. As no significant amount of insoluble protein was detected, PEI was implemented in high-scale purification. Protein was efficiently recovered from the clarified lysate ([**L**], [**Ft**] – sample before and after incubation with the chromatography, respectively). Protein in the eluate fraction [**E**] was subjected to cleavage with Ulp1 protease [**Ulp**]. After cleavage [**Cd**] and reloading the sample on Ni-agarose beads protein was ultimately polished with size-exclusion chromatography and monomeric fraction [**P2**] was separated from the dimer [**P1**]. Protein forms dimers regardless pH and ionic strength of a buffer **(B)**.

Size-exclusion chromatography indicated that TNFR1 exists in both monomeric and dimeric states. To investigate the possibility of shifting the monomer/dimer equilibrium, we decided to conduct size-exclusion chromatography in various buffers with differing ionic strength and pH, however we did not observe a substantial impact on the monomer-to-dimer ratio (Figure 3B). However, it can be observed that higher salt concentrations favor dimer formation, likely due to an increased hydrophobic effect. Ultimately, the final yield of a monomeric protein production and purification reached 6 mg/L.

### TNFR1^R347A^ stability studies

Differential scanning fluorimetry (DSF), allows for fast optimization of buffer conditions by assessment of protein stability in a thermal shift assay (TSA) [23]. Purified TNFR1^R347A^ was submitted to quality control by DSF assay. The shape of the protein melting curves in the standard formulation buffer (50 mM HEPES pH=7.4, 300 mM NaCl, 1 mM TCEP) was unsatisfactory for analysis due to lack of sigmoidal pattern of melting (**Błąd! Nie można odnaleźć źródła odwołania.**A). Therefore, a panel of 96 buffer conditions (RUBIC buffer screen, Molecular Dimensions) [24] was tested in TSA to identify optimal solution conditions. Data are presented as a heatmap with the average melting point for each condition (Figure 4B).

**Figure 4.**
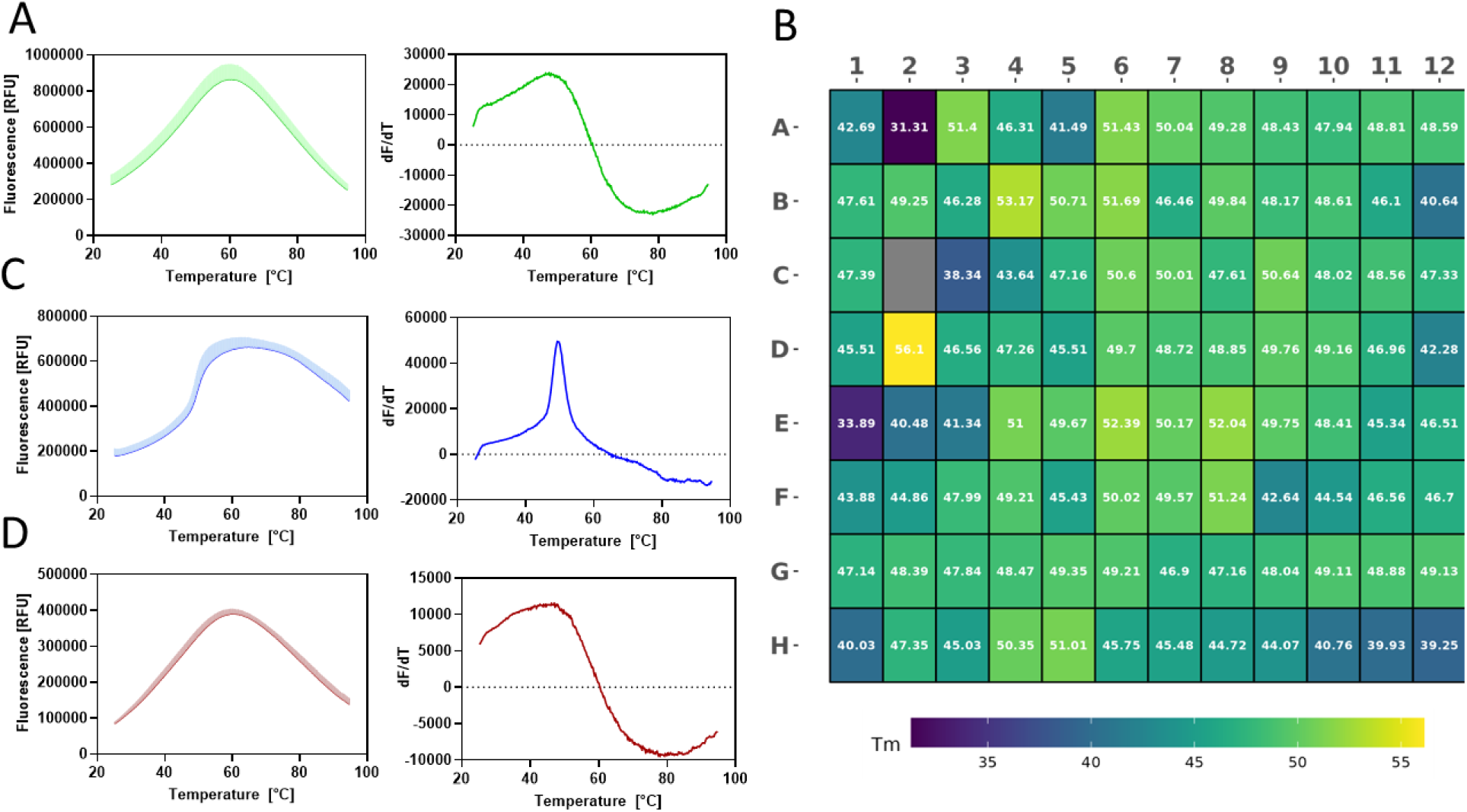
Image of data collected from DSF assay. Melting curves of the TNFR1^R347A^ in 50 mM HEPES pH=7.4, 300 mM NaCl, 1 mM TCEP (recombinant protein formulation buffer) **(A)**. Heatmap of change in melting point of protein in RUBIC buffer screen conditions. Data are presented as a mean value from two independent repeats **(B)** (Layout of the screen depicts supplementary Figure S2). TNFR1^R347A^ melting curves at pH=6.0 (SPG buffer**) (C)**. TNFR1^R347A^ melting curves at pH=9.0 (SPG buffer) **(D**).

For most of the tested conditions, the melting temperature was around 45-50°C. We noticed that the type of tested buffer has more significant impact on the shape of the curve than on the melting point. We observed that the melting curves in MES, MOPS, and Bis-Tris buffers have a sigmoidal shape, and the melting temperature determined from the first derivative is reliable. Moreover, we show that the pH value has the most significant impact on protein stability. At pH 6, we remarked on the optimal curve shape and the highest melting point (Figure 4C). However, with the increase of pH, the shape of the melting curves deteriorated, precluding precise determination of the melting temperature. In the RUBIC buffer screens the effect of salt concentration is mainly tested in the pH range of 7.5-8, at which the protein is generally unstable. Therefore, in the next experiment, we decided to check the impact of pH and ionic strength on the stability of the TNFR1^R347A^ in the MES buffer (*Supplementary*: Figure S3). We decided to quantify the impact of pH and ionic strength of the solution on protein stability. Based on the results of the DSF-guided buffer screen described above, purified TNFR1^R347A^ was thermally denatured in 0.1 M MES buffers within the pH range 5-7 and NaCl concentration between 50 and 500 mM. Data points were selected based on the Face-Centered Central Composite design. It allows the detection of not only main effects but also interactions between them and curvature effects – if present, enabling the calculation of critical points. Averaged values (technical replicates n=3) for each point were used for model training. Analysis of variance (ANOVA) conducted for model (0.1, commonly employed for this design, demonstrated that only the linear effect of pH was statistically significant. Ultimately, the model itself was not statistically significant in explaining the variance of the data (p>0.05).

**Table 1.** ANOVA results for second-order polynomial model.

|  | DF | Sum of squares | Mean square | F statistic | Prob (F statistic) |
| --- | --- | --- | --- | --- | --- |
| **NaCl** | 1 | 14.09 | 14.09 | 1.59 | 0.30 |
| **pH** | 1 | 127.36 | 127.36 | 14.36 | **0.03** |
| **NaCl:pH** | 1 | 3.14 | 3.14 | 0.35 | 0.59 |
| **NaCl<sup>2</sup>** | 1 | 0.22 | 0.22 | 0.03 | 0.88 |
| **pH<sup>2</sup>** | 1 | 5.98 | 5.98 | 0.67 | 0.47 |
| **Model** | 5 | 150.80 | 30.16 | 3.40 | 0.171 |
| **Error** | 3 | 26.60 | 8.87 |  |  |
| **Total** | 8 | 177.40 |  |  |  |

R<sup>2</sup> = 0.85; Adjusted R<sup>2</sup> = 0.6; Bayesian information criterion (BIC) = 48.48

We subsequently developed a statistically significant linear model, with homoscedastic (Shapiro p>0.05) and normally distributed residuals (Breusch–Pagan p>0.05). Nevertheless, ionic strength still proved to be insignificant within the tested range of concentrations and pH scope. The extra sum-of-squares F test confirmed that the linear model was sufficient in explaining the variance (p-value>0.05), and additional terms in the second-order polynomial model did not significantly contribute to reducing the error. To summarize, only pH exerted a significant, linear effect on protein stability in examined scope of pH and NaCl concentration values.

**Table 2.** ANOVA results for linear polynomial model.

|  | DF | Sum of squares | Mean square | F statistic | Prob (F statistic) |
| --- | --- | --- | --- | --- | --- |
| **NaCl** | 1 | 14.09 | 14.09 | 2.35 | 0.18 |
| **pH** | 1 | 127.36 | 127.36 | 21.25 | **<0.01** |
| **Model** | 2 | 141.45 | 70.72 | 11.80 | **0.01** |
| **Error** | 6 | 35.96 | 5.99 |  |  |
| **Total** | 8 | 177.40 |  |  |  |

R<sup>2</sup> = 0.79; Adjusted R<sup>2</sup> = 0.73; Bayesian information criterion (BIC) = 44.6

## Methods

### Small scale production of TNFR1^R347A^, MyD88, IRAK4 death domains in E. coli BL21(DE3) cells

Codon optimized DNA fragments encoding respective open reading frames (ORFs) (*Supplementary*: Table S1) were introduced into pET28a vectors for transformation of *Escherichia coli* BL21(DE3) bacterial strain. For each construct, a single clone was used to inoculate 10 mL of Luria-Bertani (LB) (Sigma-Aldrich, Cat. No.: L3522) broth containing 50 µg/mL kanamycin. After inoculation, bacteria were cultivated at 37 ℃ over the night (ON). The following day 15% glycerol stocks of bacteria were prepared, and seed cultures were back-diluted (1:50 ratio) to 250 mL and cultivation at 37 ℃ lasted until they reached OD_600_ 0.5 – 0.6. Next, 25 mL of cultures were transferred into 125 mL flasks and protein production was induced either with 0.1 or 1 mM Isopropyl β-D-1- thiogalactopyranoside (IPTG). Flasks were incubated with 90 rpm shaking under different conditions: (I) 37 ℃ / 3 hours, (II) 25 ℃ / ON, (III) 16 ℃ / ON.

### Small scale purification using Ni-agarose chromatography beads

After incubation under respective conditions, 2 mL of each culture was collected and cells were harvested by centrifugation (6,000 rcf; 5 min; 4 ℃) Pelleted cells were suspended in B-PER (ThermoFisher, Cat. No.: 90084), supplemented with 1 mM TCEP, cOmplete EDTA-free protease inhibitor cocktail (Roche, Cat. No.: 05056489001), 10 U/mL Viscolase (A&A Biotechnology, Cat. No.: 1010-25), 5 mM imidazole and incubated at room temperature (RT) for 20 minutes, briefly vortexing every 10 minutes. After incubation, cells were disrupted by freeze/thaw, and whole cell lysate fraction (WCL) was collected for SDS-PAGE analysis. Cell debris were separated from soluble fraction by centrifugation (20,000 rcf; 10 min; 4 ℃) and SDS-PAGE samples from supernatant were prepared (Sup). Supernatant (0.74 mL adjusted to 1 mL with lysis buffer) was incubated with Ni-agarose beads (PureCube, Cat. No.: 75103) for 1 hour at 8 ℃ with gentle mixing. Recovered chromatography beads were washed with 1 mL of 50 mM Tris/HCl, 300 mM NaCl, 5% glycerol, 1 mM TCEP, 20 mM imidazole, pH 8.0, and eluted with 0.15 mL of 50 mM Tris/HCl, 300 mM NaCl, 5% glycerol, 1 mM TCEP, 0.5 M imidazole, pH 8.0. SDS-PAGE samples from eluate (E) fractions were prepared. For SDS-PAGE analysis all samples except from eluate fractions were normalized based on OD_600_, loading 20/OD. Eluate fractions were normalized based on cell culture volume, loading cells derived from 0.2 mL of culture.

### High scale purification of TNFR1^R347A^ death domain

Previously prepared glycerol stock was used to inoculate 50 mL of LB with 50 μg/mL kanamycin. Bacteria were cultivated over the night at 37 ℃ with 220 rpm aeration to generate seed culture. The following day the seed culture was used to inoculate 2 L of LB with 50 μg/mL kanamycin (1:50 dilution ratio). Cells were further cultivated with 180 rpm aeration at 37 ℃ until they reached OD_600_ 0.53. Protein production was induced with 0.1 mM IPTG, and flasks were incubated at 25 ℃ over the night with 90 rpm aeration. Afterwards cells were harvested by centrifugation (6,000 rcf; 20 min; 4 ℃) and suspended in ice-cold 50 mM Tris/HCl, 0.3 M NaCl, 5% glycerol, 5 mM imidazole, 10 mM β-mercaptoethanol (BME), cOmplete EDTA-free protease inhibitor cocktail (Roche, Cat. No.: 05056489001), 5 U/mL Viscolase (A&A Biotechnology, Cat. No.: 1010-25), pH 8.0 and passed twice through an EmulsiFlex at 15,000 psi. Afterwards 5% polyethylyneimine (PEI) was added to the homogenate reaching 0.1% [21], and cell lysate was centrifuged (20,000 rcf, 30 min, 4°C) to sediment the insoluble material. Protein was recovered from bulk of host cell proteins by immobilized-metal affinity chromatography (IMAC) using 10 mL of Ni-agarose beads (PureCube, Cat. No.: 75103). After 1 hour of incubation at 8 °C with gentle mixing, Ni-resin was collected and washed with 50 mM Tris/HCl, 0.3 M NaCl, 5% glycerol, 5 mM BME, 20 mM imidazole, pH 8.0. Protein was eluted with 40 mL of 50 mM Tris/HCl, 0.3 M NaCl, 5% glycerol, 5 mM BME, 0.5 M imidazole, pH 8.0 and in the presence of his-tagged Ulp1 protease (TNFR1- DD: Ulp1 mass ratio 100: 1), sample was dialyzed over the night against 1 L of the same buffer but without imidazole. After centrifugation dialysate was reloaded on Ni-agarose beads, and flow-through after concentration was ultimately polished by size-exclusion chromatography using HiLoad Superdex 26/600 75 pg column (Cytiva) equilibrated with 50 mM HEPES/NaOH, 0.3 M NaCl, 1 mM TCEP, pH 7.4. Protein was used in Differential Scanning Fluorimetry (DSF) analysis.

### Differential Scanning Fluorimetry (DSF)

Differential scanning fluorimetry was performed using ViiA 7 Real-Time PCR System (Applied Biosystems). All measurements were performed on a 384-well PCR plate (4TITUDE, Cat. No. 4ti-0387). The final protein concentration in all experiments was 10 µM. 10x SYPRO Orange (Invitrogen^TM^, Cat. No. S6650) was used as a dye and it was dispensed by Echo 555 liquid handler (Labcyte Inc.). The assay volume was 10 µl and the final DMSO concentration in the assay was 1%. In the buffer screen experiment 7.4 µl of commercial RUBIC Buffer Screen (Molecular Dimensions, Cat. No. MD1-96) was used. The experiment on the dependence of stability on pH function and ionic strength was conducted in 0.1 M MES with different ranges of pH (5-7) and ionic strength (50-500 mM). The plates were sealed, mixed and centrifuged. DSF measurements were taken at excitation 470 nm to emission 623 nm with the temperature range 25°C- 95°C and ramp rate 0.1°C/s. The melting point was calculated by the first derivative using Protein Thermal Shift Software (Applied Biosystems).

### Statistics and data interpretation

Face-Centered Central Composite for 2 factors (*f*) was applied for the preparation of an experimental design (Table 3). It comprises of a full factorial design (2^*f*^) with additional star points (2*f*) and a center point. In total 9 points were used and each point was technically replicated in the DSF experiment.

**Table 3.**
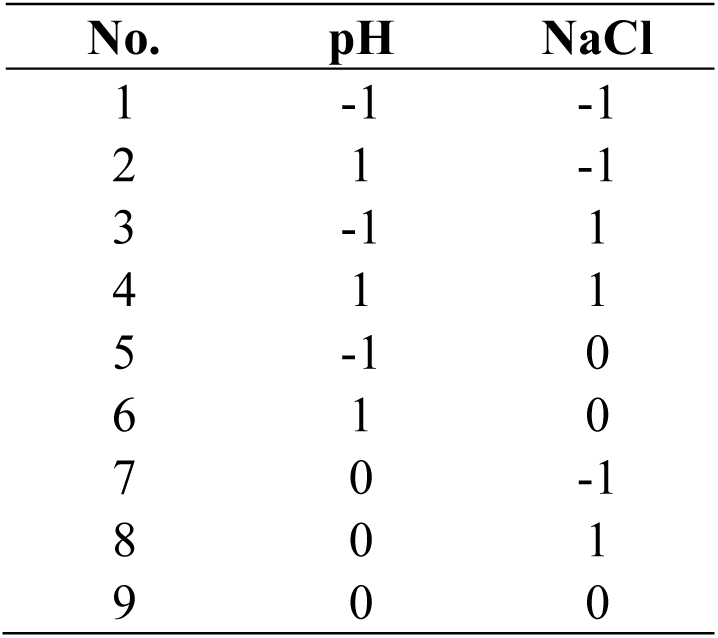
Face-Centered Central Composite Design for NaCl, pH factors with 1 center point. Numbers - 1 and 1 denotes the maximal and minimal values.

For describing the relation between a response and considered factors *f*, a second-order polynomial model (Equation (0.1) is ubiquitous in response surface designs [25]:

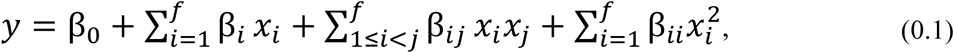

where y is the response, β_0_ the intercept, β_i_ the main coefficients, β_ij_ the two-factor interaction coefficients and β_ii_ the quadratic coefficients.

Data obtained from the DoE was analyzed using open-source libraries: Statsmodels [26], Scipy [27] and Pandas [28] compatible with Python 3.11

